# Connecting people and ideas from around the world: global innovation platforms for next-generation ecology and beyond

**DOI:** 10.1101/012666

**Authors:** Peter Søgaard Jørgensen, Frederic Barraquand, Vincent Bonhomme, Timothy J. Curran, Ellen Cieraad, Thomas G. Ezard, Laureano A. Gherardi, R. Andrew Hayes, Timothee Poisot, Roberto Salguero-Gómez, Lucía DeSoto, Brian Swartz, Jennifer M. Talbot, Brian Wee, Naupaka Zimmerman

## Abstract

We present a case for using global community innovation platforms (GCIPs), an approach to improve innovation and knowledge exchange in international scientific communities through a common and open online infrastructure. We highlight the value of GCIPs by focusing on recent efforts targeting the ecological sciences, where GCIPs are of high relevance given the urgent need for interdisciplinary, geographical, and cross-sector collaboration to cope with growing challenges to the environment as well as the scientific community itself. Amidst the emergence of new international institutions, organizations, and dedicated meetings, GCIPs provide a stable international infrastructure for rapid and long-term coordination that can be accessed by any individual. The accessibility can be particularly important for researchers early in their careers. Recent examples of early career GCIPs complement an array of existing options for early career scientists to improve skill sets, increase academic and social impact, and broaden career opportunities. In particular, we provide a number of examples of existing early career initiatives that incorporate elements from the GCIPs approach, and highlight an in-depth case study from the ecological sciences: the International Network of Next-Generation Ecologists (INNGE), initiated in 2010 with support from the International Association for Ecology and 18 member institutions from six continents.

---

Scientists have entered a new age of research where excellence and impact are driven by international collaboration (Adams 2013). Although there is much focus on the improved quality of research outputs (Parker et al. 2010), international collaboration may also increase the quality of a broad set of basic practices in the scientific community. Ultimately, these practices influence the quality of research, and of policy, educational, and communication outputs. Online communication tools that facilitate a common community infrastructure are essential for international collaboration. The benefit of online infrastructures may be particularly significant in the modern era, helping to ameliorate the difficulties inherent in connecting a growing global research community across geographical space, disciplines and research questions. The global community of ecological scientists, defined as including researchers studying all applied and basic aspects of ecology and evolution, stands to benefit from these growing interconnections as it is increasingly made up of researchers from diverse backgrounds (Carpenter et al. 2009), a trend that has been building for decades (Sugden 1986).

An online infrastructure allows a community to pool fragmented resources, more rapidly exchange knowledge sources, and develop and share new communal assets. Such assets include but are not limited to: (1) training on new and emerging issues (Walker 2013), (2) framing new approaches to the practice of science (Molloy 2011), (3) preserving knowledge and data that would otherwise be lost (Reichman et al. 2011, Fox 2012), (4) establishing a common voice on international issues (Beaudry et al. 2010, Alberts 2011, Future Earth 2013, Mooney et al. 2013, Griggs et al. 2013), and (5) facilitating information gathering about the community itself (Parker et al. 2010, Barraquand et al. 2014). Locally, insufficient resources often limit the development of these community assets. However, significant contributions can be made by pooling otherwise disparate efforts from a global community into a shared common space that is openly accessible.

In addition to helping develop resources in communities, a shared infrastructure provides a platform from which to address a broad suite of challenges. For example, one widespread methodological challenge is encouraging more transparent research practices (Evans and Reimer 2009, Voronin et al. 2011, Boulton et al. 2011, Enserink 2012, Mace 2013), without inadvertently threatening scientific integrity (e.g. through the emergence of predatory journals (Bohannon 2013)). Another challenge is to increase participation of underrepresented groups within the scientific community—a change that is crucial for promoting public interest in science and for improving scientific practice overall (Hampton et al. 2013). Finally, there is growing awareness of a mismatch between the supply of new scientists and the availability of jobs for which they have been trained (Stephan 2012). A more inclusive international discussion may help to address these challenges more rapidly.

In this Innovative Viewpoint, we present a case for one approach to international collaboration via online infrastructures. We describe a particular type of online infrastructure, denoted here as Global Community Innovation Platforms (GCIPs). GCIPs strive to engage large parts of the international community through a relatively flat and open organizational structure that ensures many individuals can contribute to the innovation process. We are especially concerned with the potential benefits of GCIPs for the growing communities of early career scientists. To better illustrate what differentiates GCIPs from other, existing, efforts in the scientific community, we provide examples of organizations that fall within this conceptual framework and highlight the benefits they provide. Throughout, we complement such examples with a recent case study from the ecological sciences, the International Network of Next-Generation Ecologists (INNGE) (Jørgensen et al. 2011).

## Defining GCIPs

Global Community Innovation Platforms exhibit four main features. These include a commitment to: (1) global reach and membership; (2) improving connections between local communities; (3) exploring and promoting new and emerging paradigms; and (4) being an accessible repository of information for and about the community (Table 1). Although relatively few initiatives contain all four characteristics, we here provide examples of networks and organizations that currently incorporate one or more of these features.

**Table 1.**
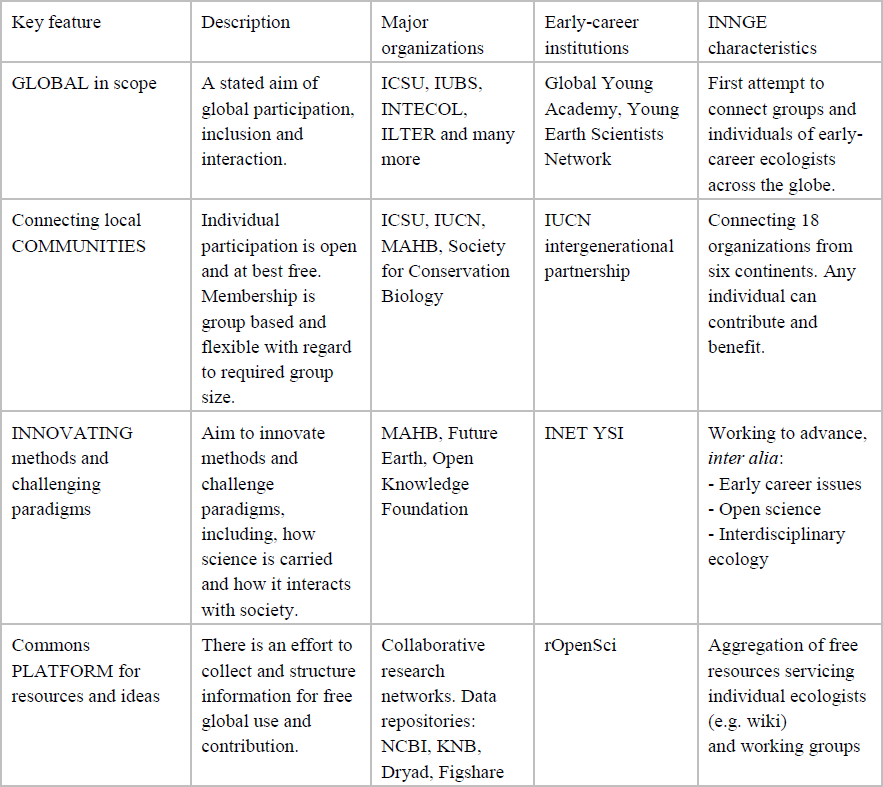
Four proposed defining features for Global Community Innovation Platforms. Examples of institutions with resembling characteristics are given for each feature, including early career institutions. Characteristics of INNGE are highlighted.

*(1) Global reach and inclusivity:* GCIPs have a specific aim of a global constituency and effect a strategy to reach and maintain this aim. Although there are a large number of international scientific organizations, including the International Council for Science (ICSU) and the International Social Science Council (ISSC), examples that target researchers early in their careers are uncommon (but consider, for example, the Global Young Academy, an international network of excellent young scientists (Beaudry et al. 2010, Alberts 2011)). This global aim is in contrast to many scientific organizations (e.g. national learned and professional societies) that are highly effective in serving their members, but are not explicitly international in scope. GCIPs do not compete with these groups, but can help to link them together. This broad scope brings particular challenges as well: the successful maintenance of an international organization in the long-term can depend on stable financial resources and secretariat assistance, both of which are often localized to one or more locations in the short term.

*(2) Connecting and servicing communities:* GCIPs aim to connect existing national or local-scale communities and to support the development of such infrastructures where they might be absent. Implementation of most on-the-ground initiatives can only be carried out at the local scale and localized infrastructures are therefore a precondition for many ideas to go from paper to practice. Strengthening of local groups ensure that ideas conceived or piloted in one place can be put into practice elsewhere. Many networks and associations share this goal, including the International Association for Ecology (INTECOL), the International Union for Conservation of Nature (IUCN), ICSU, the Millennium Alliance for Humanity and the Biosphere (MAHB), and the Global Young Academy. In addition, international learned societies, such as the Society for Conservation Biology, offer a common ‘baseline’ infrastructure for the establishment of local chapters (Table 1).

*(3) Innovation:* GCIPs explicitly encourage the exploration of new approaches to existing challenges. Many existing groups serve this role implicitly. Two of the largest ecological societies, the British Ecological Society (BES) and the Ecological Society of America (ESAmerica), support such approaches; examples include the Open Science section of ESAmerica and the annual ecology-squared symposium of BES, which every year explores an emerging topic in ecology. For other groups, it is explicitly part of their main mission. Such examples include The Open Knowledge Foundation and others devoted to increasing transparency in science (Molloy 2011). Early career initiatives include the IUCN’s iAct dialogues, which emphasize an action-oriented approach to sustainability, and the Institute for New Economic Thinking-Young Scholars Initiative, which aims to counter a loss of diversity in economics research and teaching (Colander 2005).

(4) *Preserving and sharing information:* The fourth feature of a GCIP is the preservation and dissemination of community knowledge assets. These assets—which may include ideas, data, tools, or methodologies—may be generated within or external to the GCIP. The fourth feature can be sought via open online platforms that any individual can contribute to. Open online platforms lower the barriers to broader participation, such as those imposed by financial costs memberships and logistics costs. Research networks and data repositories already use these infrastructures. Distributed research networks often act as online platforms that share information from geographically replicated research projects. International examples include the ILTER network as well as newer initiatives such as Drought-Net, NutNet, and PlantPopNet. There are additionally many examples of repositories that archive and make research data available - a comprehensive list is available at the online Registry of Research Data Repositories. Building beyond research coordination and data accessibility, the “open science” movement advocates for transparency and sharing throughout the research process (Hampton et al. 2014). Numerous groups develop software tools and provide resources to the broader community with the aim of end-to-end reproducibility (Mascarelli 2014). An example is the developer collective rOpenSci, which has a strong early career component. GCIPs include and expand the above uses of online platforms to the many other activities in the scientific community beyond research itself. For example, GCIPs help disseminate opportunities for funding and employment, share success stories, and promote upcoming events. These can be shared via mailing lists, online debate fora, micro-blogging platforms such as twitter (Bik and Goldstein 2013), and knowledge sharing structures such as wikis (Wikipedia being the best-known example in the general public) (Mascarelli 2014).

## Benefits of an early career GCIP for the ecological sciences

The ecological sciences are in need of initiatives that strengthen interdisciplinary, geographical and cross-sector collaboration (Lubchenco 1998, Mace 2013). Opportunities to develop these initiatives abound: there are many new international institutions and organizations (Kaplan 2013), a growing number of national and international meetings, and a burgeoning wealth of new information communicated online and via social media (Bik and Goldstein 2013). In addition to the value of addressing these needs, a GCIP in the ecological sciences could provide particular value to early career ecologists. A rapidly growing demographic group, early career scientists face the increasing competition for jobs (Kitazawa and Zhou 2011, Stephan 2012), a situation compounded by doctoral programs that continue to emphasize academic careers at the expense of providing training for non-academic career paths (Sauermann and Roach 2012, Colebunders 2012). GCIPs fill a particular niche by providing opportunities for long-term, rapid and coordinated international interaction by early career scientists.

Within this context, PhD students and postdoctoral researchers from ecological societies around the world founded the International Network for Next-Generation Ecologists (INNGE) in 2010 with crucial support from the International Association for Ecology (INTECOL). Today, INNGE connects 18 organizations from six continents, many of which are ecological societies (Fig. 1). These include large societies such as the British Ecological Society, the Ecological Society of America and the Ecological Society of Australia as well as international societies, such as the International Biogeography Society (complete list here). Combined, these members represent thousands of early career ecological scientists. INNGE has focused on creating opportunities for early career scientists to carry out collaborative and innovative activities through global communication and outreach (Box 1). INNGE also aims to facilitate contributions from the entire scientific community on emerging topics, such as the interconnected challenges associated with achieving long-term sustainability of current human activities, evidence-based reform of university curricula, and increased adoption of open science practices. INNGE works primarily via online communication tools that can be contributed to and accessed by any person, these include listservs, shared task management systems (e.g. Slack and Trello), Voice over IP services (e.g. Skype and Fuze), and a wiki (http://innge.net/wiki). These tools enable a broad reach: INNGE has had over 10,500 unique visitors to its website from 147 different countries, over 1,500 followers on Twitter, and currently reaches almost 400 members on a general mailing list and nearly 650 on a newsletter list. In the following we focus on the types of contributions early career organizations with GCIP-like characters provide to the international community, including INNGE’s recent and ongoing activities (Box 1).

**Figure 1.**
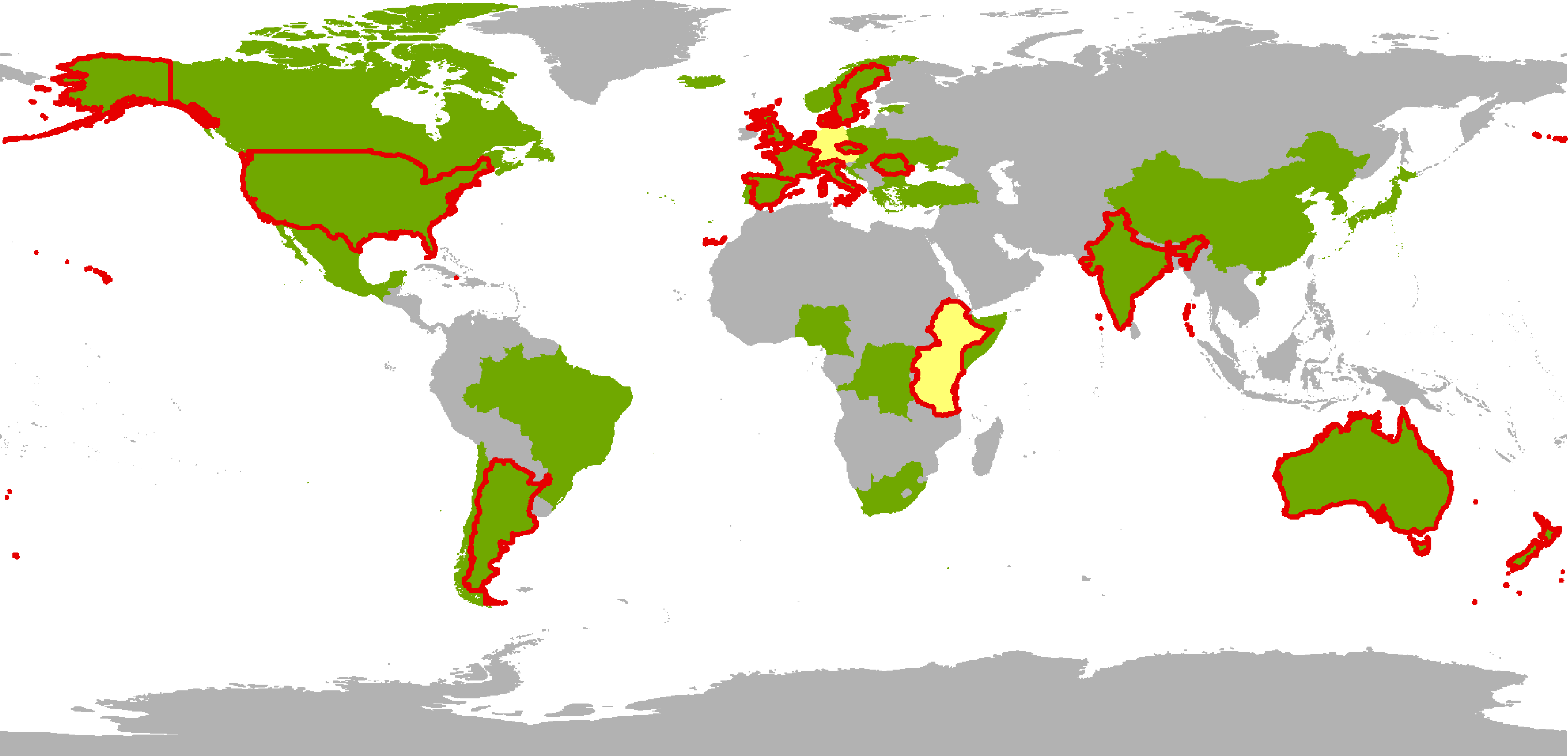
Geographical location of current national ecological societies in green and regional societies in yellow. Ecological societies from countries outlined in red are institutional members of INNGE (http://www.innge.net/node/339). Additional institutional members are the International Biogeography Society, the Millennium Alliance for Humanity and the Biosphere, and the Society for Human Ecology.

#### How has INNGE helped early-career ecologists and what collaborations have INNGE engaged in to this end?

*Examples of services that INNGE provides for early-career ecologists*

INNGE organizes **early-career events at meetings** such as the quadrennial INTECOL congress ((www.intecol2013.org)

INNGE helped initiate a **peer review mentoring scheme** in the *New Zealand Journal of Ecology* (http://newzealandecology.org/nzie/nzie-reviewer-mentoring-scheme) (Curran et al. 2013)(Curran et al. 2013)

INNGE provides a **forum for debate** of emerging and influential topics in ecology (http://www.innge.net/blog.http://www.innge.net/node/9)

INNGE provides advice and support to new local and regional early-career groups, such as the effort to establish a **regional network in Africa**

INNGE provides **surveys of early career opinion** (Barraquand et al. 2014)

INNGE hosts EcoBloggers, an open aggregator of more than 70 ecology blogs (http://www.innge.net/ecobloggers)

*What initiatives has INNGE worked with?*

INNGE is in close collaboration with **INTECOL** (http://intecol.org) and represented on INTECOL’s executive board and the local organizing committee of the INTECOL congress.

INNGE is supported through the membership of **18 member institutions** (http://www.innge.net/node/339)

INNGE has collaborated with open science initiatives such as **F1000 Research, Peerage of Science and Scholastica** through interviews communicating their work and mission (http://www.innge.net/blog)

INNGE is collaborating with the **International Council for Science** (**ICSU**, http://www.icsu.org/) and the **International Social Science Council** (**ISSC**, http://www.worldsocialscience.org/) to provide early career opportunities in Future Earth (http://www.icsu.org/future-earth). This includes a **Young Scientists Networking Conference on Integrated Science** held in Italy in May 2014.

INNGE is collaborating with the **Global Young Academy** (http://www.globalyoungacademy.net/) and members of the **Young Academy of Sweden** and **Young Academy of Germany** to raise awareness about early career issues in **Future Earth**

INNGE is collaborating with the **Institute for New Economic Thinking’s Young Scholars Initiative (INET YSI**,http://ineteconomics.org/ysi) to organize events that focus on building better interdisciplinary bridges between ecology and economics.

INNGE participated as early-career stakeholders in **BiodiversityKnowledge** an EU funded research project that explores the opportunities for establishing a European science-policy interface on biodiversity and ecosystem services.

### A hub of early career knowledge

An important and basic function of any GCIP is to work as a knowledge hub that individuals and local groups can tap into for advice and information. For example, organizations, such as The Early Career Climate Forum (ECCF) support an online forum where researchers and professionals can share information about resources, ideas, and projects related to climate change research. The Young Earth Scientist (YES) Network, an international network for early career geoscientists that aims to promote earth science for the benefit of society, publishes a newsletter highlighting opportunities for network members. INNGE’s wiki (which anyone can edit after registering) serves as such a hub, giving an overview about everything from useful email listservs to open access publishing options. In addition, INNGE devotes a part of its own blog to publishing Next-Generation Point-of-View posts, which feature ideas identified by the community as particularly interesting. Although some posts highlight papers written by early career scientists, such as how to prepare for a career outside academia (Blickley et al. 2013), others have developed from an initial post into a peer-reviewed publication (Desjardins-Proulx et al. 2013). A recent post describes an example of successful educational outreach to children from lower income families initiated by an early career ecologist (the University of Arizona’s Sky School), an initiative that in 2014 was recognized by the U.S. federal government.

Knowledge sharing is as important among and between organized early career groups as it is between individual researchers. A central challenge for many early career groups is building a critical mass of individuals that persists despite organizational turnover. The lack of available best practices for starting and maintaining such groups set new groups at a disadvantage. GCIPs can help: the YES Network offers resources for national chapters, and INNGE’s website features a showcase section where early career groups can highlight their past and ongoing activities. The latter serve to inspire other groups facing similar challenges. In one example, the Ecological Society of America’s Student Section worked closely with the ESA’s executive board after the 2010 Gulf of Mexico oil spill to organize a database of Gulf Coast ecological data to serve as a baseline for post-spill comparisons (Ramos et al. 2012).

### Distributor of online debates

Even in a well-connected world, online debates can emerge as regionally structured (Ardichvili et al. 2006, Pfeil et al. 2006, Jiang et al. 2009). Such regional structure may lead to inefficient transfer of ideas across larger geographic or cultural contexts. For example, although science blogs can stimulate debate (Fox 2012), their reach can be compromised by the time it takes to monitor the jungle of individual blogs as they come and go (Bonetta 2007, Wilkins 2008). Thus, there is value in stable points of aggregation. A blog aggregator works by collecting the posts from a set of contributing sites and turning them into a single feed (Wilkins 2008). As all interested bloggers can contribute, readers benefit from exposure to a more diverse set of viewpoints. Bloggers can in turn increase their own readership, which is especially useful when establishing a new blog. A blog aggregator is an example of how GCIPs can help enhance the online voice of individuals and foster a sense of online community. In 2013, INNGE launched *EcoBloggers*, which gathers posts from more than 80 blogs and blogging communities. It was heavily inspired by *r-bloggers*, which gathers posts on the statistical programming language R (R Development Core Team 2010). Aggregators encourage readers to consider everything from the thoughts of undergraduate students to posts by editors of highly regarded journals.

### Surveying the community

Because of their globally distributed and online nature, GCIPs are especially well suited to survey international communities. International surveys provide means to inform the debate on what would otherwise be national issues, such as curricula reform in higher education (Salguero-Gómez et al. 2008, Zimmerman et al. 2009). For example, the YES Network conducted one survey on the unique challenges women face as geoscientists, and another assessing the impact major career moves on the lives of geoscientists. These surveys can be distributed through coordinated action across many regional listservs. In this manner, INNGE surveyed ecologists’ experience and self-perceived skills in mathematics, statistics, and programming. Relayed through the major listservs in Europe and North America, including ECODIFF and ECOLOG-L, within a week the survey had close to 1,000 respondents. The results of the survey demonstrated a dissatisfaction with the level of mathematical training that ecologists receive during their formal education, a dissatisfaction that was consistent across continents (Barraquand et al. 2014). Such surveys illustrate the ability of GCIPs to provide evidence-based suggestions to help address the current concerns of the community.

### Promoting early career interdisciplinarity

Developing greater interdisciplinary fluency is a crucial step for enhancing the contribution that ecological researchers can make towards addressing major societal challenges (Lubchenco 1998, Liu et al. 2007, Collins et al. 2011, Bettencourt and Kaur 2011, Kueffer et al. 2012, Mace 2013). By providing information on and access to researchers in other disciplines, GCIPs can highlight paths into interdisciplinary work. For example, the Association of Polar Early Career Scientists aggregates websites that connect researchers across disciplines who are actively studying or interested in studying the same polar systems. The Intergenerational Partnership for Sustainability has documented the goals and achievements of their sustainability groups comprised of researchers with different scientific backgrounds from over 40 different countries. INNGE has been engaged in three concrete initiatives to promote interdisciplinary exchange. In the first, INNGE collaborated with the Institute for New Economic Thinking -Young Scholars Initiative (INET YSI) to showcase interdisciplinary dialog between ecology and economics. Inspired by a series of annual summits that bring together senior ecologists and economists to discuss important societal issues (Söderqvist et al. 2011), the first project from this collaboration was a free online webinar series featuring senior scientists working across the boundaries of ecology and economics (Box 1). The second initiative, in conjunction with the ten-year Future Earth initiative (Future Earth 2013), produced concrete suggestions to help realize Future Earth’s goal of “engaging a new generation of young scientists”. These recommendations were developed with both the Global Young Academy and national young academies of scientists, establishing an organized voice for early-career scientists within Future Earth (Box 1). Finally, building on the first two initiatives, INNGE, INET YSI, ICSU, and ISSC, with funding from the German Research Foundation (DFG), co-organized the 2^nd^ Future Earth young scientist networking conference in 2014. The conference brought nearly 30 young scientists from around the world together to advance interdisciplinary approaches on the topic of “ecosystems and human well-being in the green economy”. Several collaborative research projects emerged from this meeting.

## How to get involved in an early-career GCIP

GCIPs offer novel opportunities for scientists to increase their influence and the success of GCIPs rely on engaging diverse and creative groups of scientists. So, how does one get involved in a (early career) GCIP?

*1) Find out what you like doing.* Almost everyone has something that they like doing that can be of benefit to others. No matter to what extent you decide to become involved in a GCIP, your efforts will be most useful if you focus on a topic that brings you personal satisfaction. Such efforts could include contributing a blog post, developing online infrastructure, fundraising, facilitating skill building, doing data analysis, arranging meetings, and coordinating interdisciplinary projects—or anything else you can dream up!

*2) Show-up.* In many GCIPs, you can interact with people simply by sending an email to a listserv or commenting on a website. However, to become more involved, get in touch with the organization’s leaders or coordinators of sub-groups. If you are attending a meeting, consider e-mailing in advance to inquire about discussing opportunities in person. If an in-person meeting is difficult, consider requesting a quick Skype or phone call. In conversation, do not be afraid to highlight what you can bring to the group, but also consider inquiring about what is currently needed. Being honest about your interests, available time, energy, and preferred mode of interaction will set a good precedent for communication with the team.

*3) Stick around.* Working in groups, while providing an opportunity to contribute to larger initiatives, also requires patience. Being involved with a GCIP means coordinating workflows and staying up to date with ongoing projects. Compared to an individual research project, coordination—across many countries, time zones, and career stages—becomes a bigger part of the process. Being prepared to stay involved over time enhances the chance that you will be able to contribute and to see the results of your efforts make an impact. The above examples highlight what can be done by GCIPs at an international scale within the relatively short period of a couple of years. If you decide to be involved in a GCIP, you will not only have had a good chance to see the group evolve and grow its impact, you are also likely to have made new colleagues and friends for the future.

## Long-term viability: eliminating barriers and increasing resilience

The persistence of a long-term innovation platform may at first appear to be at odds with the rapidly changing lives and global nomadism of many early career scientists (van Noorden 2012). One challenge, for example, is that the ability to engage in active GCIP participation can change on short notice. Maximizing the resilience of early career GCIPs over the long-term is therefore a necessary consideration. INNGE’s resilience strategy is based on a backbone of institutional memberships, where groups of early career representatives succeed each other continuously as individuals move on. A central purpose of this strategy is to reduce the barrier to participation. As institutional memberships provide stability over the long term, individual ecologists are freed to realize innovative ideas. In 2014, the membership of INNGE elected its first governing board, composed of representatives spanning the six populated continents. Term limits (board members may serve a maximum of two 2 year terms) and overlapping turnover of elected positions (up to 50% turnover per year) are a second element of the resilience strategy, which aim to secure a continuous influx of new perspectives and avoid abrupt losses of institutional experience, respectively.

What activities will early career GCIPs carry out in the future? This is hard to say; innovation is an emergent group process and historical precedents are not always useful. INNGE was established to better connect the global community of early career ecologists. In doing so, it aims to enhance international knowledge transfer, and thereby foster and communicate across cultural and geographic boundaries. One hundred years after the formal establishing of the first ecological societies (the British Ecological Society in 1913 and the Ecological Society of America in 1914), the field of ecology is faced with an extraordinary challenge: to elucidate the planetary boundaries for human activities (Rockström et al. 2009a, 2009b) and inform the global governance of the environments rapidly being altered by our species (Mace 2013). In the search for sustainable solutions that are grounded in the natural and social sciences (Burger et al. 2012), geographical, institutional, and disciplinary barriers in the scientific landscape must be less predominant. GCIPs provide one way by which communities can get together to help break down these barriers for the benefit of their disciplines and the generations that follow. Beyond the purpose of a GCIP in ecology, we advocate that GCIPs are of general use to speed innovation in distributed communities of scientists. We hope that the example of INNGE will further stimulate the development of GCIPs in other scientific fields.

## Acknowledgments

We thank all of those people from around the world who have contributed to INNGE for inspiring this manuscript. Alan Covich, Georgina Mace, and Harold Mooney provided insightful comments that improved a previous version of this manuscript. We thank the INTECOL and the British Ecological Society and the Ecological Society of America for their continued interest in and support of INNGE’s activities.

